# Inherently confinable split-drive systems in *Drosophila*

**DOI:** 10.1101/2020.09.03.282079

**Authors:** Gerard Terradas, Anna B. Buchman, Jared B. Bennett, Isaiah Shriner, John M. Marshall, Omar S. Akbari, Ethan Bier

## Abstract

CRISPR-based gene drive systems, which copy themselves based on gene conversion mediated by the homology directed repair (HDR) pathway, have potential to revolutionize vector control. However, mutant alleles generated by the competing non-homologous end-joining (NHEJ) pathway that are rendered resistant to Cas9 cleavage can interrupt the spread of genedrive elements. We hypothesized that drives targeting genes essential for viability or reproduction also carrying recoded sequences to restore endogenous gene functionality should benefit from dominantly-acting maternal clearance of NHEJ alleles, combined with recessive Mendelian processes. Here, we test split gene-drive (sGD) systems in *Drosophila melanogaster* that were inserted into essential genes required for viability (rab5, rab11, prosalpha2) or fertility (spo11). In single generation crosses, sGDs copy with variable efficiencies and display sex-biased transmission. In multi-generational cage trials, sGD follow distinct drive trajectories reflecting their differential tendencies to induce target chromosome damage or lethal/sterile mosaic phenotypes, leading to inherently confineable drive outcomes.

## Introduction

There is excitement, particularly in entomological fields, regarding the potential use of CRISPR-based gene drive systems to control pest populations [1] or disease-transmitting vectors [2–5]. Genetic engineering of populations ideally results in complete introgression of the desired beneficial genetic element into native backgrounds with minimal release of transgenic individuals, which can be achieved with efficient “low-threshold” gene-drive systems (capable of spreading from a low seeding frequencies) such as CRISPR-Cas9 drives. These drive systems rely on a CRISPR-based cut-and-repair mechanism, wherein Cas9 produces a double-stranded break (DSB) in the DNA followed by homology directed repair (HDR) using the homologous chromosome as a template. This results in copying of the genetic element into the break on the Cas9-cleaved chromosome [6, 7]. However, generation of alleles resistant to Cas9 cleavage [8, 9] via the competing non-homologous end-joining (NHEJ) pathway can interrupt the spread of the desired genetic element [10], particularly if the cleavage-resistant allele carries less of a fitness cost than the driving allele. NHEJ-induced alleles typically consist of short insertions or deletions (indels) that result in amino acid substitutions or frameshifts that modify the gRNA target sequence, preventing further Cas9 cleavage [11]. Also, non-functional drive-resistant alleles, which create loss-of-function (lof) mutations and are typically the most abundant NHEJ products, can delay complete drive introgression due to their being only gradually eliminated by negative selection when homozygous [10]. Therefore, overcoming the generation and/or persistence of such cleavage resistant alleles is essential to the long-term success of CRISPR-based drives in the field.

Several strategies have been proposed to reduce the incidence or effects of cleavage-resistant alleles. One approach is to employ tightly regulated germline-specific promoters to avoid somatic expression of Cas9 that can lead to NHEJ-induced repair events [9, 10, 12]. Some germline promoters are quite specific, at least in certain genomic contexts (e.g., nanos [13] or zpg [14]), while others are leaky (e.g., vasa promoter [15, 16]). Another tactic is to identify highly conserved genomic target sites as cleavage sites such that all NHEJ mutants suffer severe fitness costs [14, 17, 18]. Multiplexing gRNAs have also been suggested as a potential means for reducing the impact of resistance [19], but has only shown modest benefits at best [12, 20].

Recently, it has been suggested that drives targeting conserved genes essential for survival or fertility that also carry a recoded cDNA restoring endogenous gene activity would benefit from two forms of positive selection in populations [2, 17, 21, 22]. The first advantage results from gradual Mendelian culling of recessive deleterious NHEJ alleles. The second, more rapid, depends on an active dominant process referred to as lethal/sterile mosaicism, in which maternal deposition of Cas9/gRNA complexes into the embryo mutates the paternal allele in a mosaic fashion [23]. If progeny inherit the recoded drive, they are protected from this maternal effect since they carry one unassailable functional allele. Individuals inheriting a non-functional NHEJ allele, however, will lack gene activity in many cells if large-scale mosaicism occurs and the targeted gene is required broadly in the organism for either viability or fertility. Thus, lethal/sterile mosaicism acts dominantly to generate (and eliminate) NHEJ alleles as they arise and acts in concert with standard negative selection against recessive deleterious alleles, the latter leading to their steady decay.

Two main types of CRISPR-based gene drives, or “active genetic” elements, have been designed and tested: full gene drives (fGD) [2, 3, 6, 15], in which linked sources of Cas9 and gRNA inserted together at a single genomic site spread through the population, and split gene drives (sGD) [5, 16, 24, 25], where a gRNA-only cassette capable of copying via directional gene conversion is inserted in a specific genomic position and is combined with an independent Mendelian or “static” source of Cas9, located at a second locus. It is also possible for the gRNA-carrying element to include a second gRNA that cuts the genome where the Cas9 source is inserted, empowering that latter element to copy as well. In such trans-complementing systems [26], each element behaves according to standard rules of Mendelian inheritance when propagated separately, but once combined, they copy as a coupled full-drive system. In all of these cases, the gene-drive carried on one chromosome converts the homolog in heterozygous germline cells, creating a strong bias for Super-Mendelian (>50%) transmission of the drive allele. Since the gRNA containing locus in split-drive systems is the only one displaying Super-Mendelian inheritance, this method is more amenable to experimental analysis of different parameters influencing drive efficiency (e.g., different Cas9 sources can be tested in a combinatorial fashion with various gRNA-based drives) and has been deemed safer for laboratory and more ammenable for localized release purposes [5, 24, 27, 28].

In this study, we design and test several sGDs in *Drosophila melanogaster* with various genetic parameters and strategies either to limit or extend sGD drive potential. We demonstrate how resistance can be largely overcome if the drive allele carries an efficient gRNA and recoded target gene sequence. To achieve this, we inserted sGDs into target genes known to be essential for survival or fertility. sGDs were efficiently transmitted through females and to a lesser degree through males when combined with *vasa-Cas9* or *nanos-Cas9* sources located on different chromosomes. Genomic context (Cas9 promoter, chromosomal location) also contributed to gene-drive efficiency and rates of NHEJ mutagenesis. In multigenerational cage studies, sGDs carrying recoded cDNA sequences that restore target gene function upon copying benefited from the dual effect of a dominant maternal phenomenon (lethal/sterile-mosaicism) that rapidly eliminates non-functional NHEJ alleles, acting together with classic gradual zygotic Mendelian negative selection. These benefits are amplified in the case of sGDs inserted into loci demonstrating modest haplo-insufficient phenotypes. In this case, they generate naturally self-limiting and inherently confinable drive systems by virtue of imposing a fitness cost on the Cas9 transgene when co-inherited with the drive element.

## Results

### Design of split gene drive (sGD) elements in autosomal recessive lethal loci

We tested whether Super-Mendelian transmission efficiencies could be achieved in a split genedrive system inserted into essential gene targets, we engineered genetic elements that target four different loci in *D. melanogaster*, required either for viability or fertility. These elements, referred to hereafter as split gene-drives (sGDs), were crossed to different sources of germline-expressed Cas9. All sGD cassettes contain a gRNA targeting an exonic region of the gene of interest; a recoded 3’ cDNA portion of that target gene, which restores function of the locus; a 3xP3-tdTomato dominant marker, and a partial sequence of an Opie2-eGFP transgene (designed for future studies to permit conversion of each sGD into a full gene drive by hacking [29]). All the aforementioned cargo was flanked by 1kb homology arms to the gRNA cut site to support HDR-mediated integration of the cassette into the genome (FIG. 1a). When the sGD and Cas9 are carried in separate strains, they are inherited at Mendelian frequencies. However, when combined with a source of Cas9, the sGD cassette copies to the homologous chromosome and is transmitted at super-Mendelian rates (Fig. 1b). As mentioned previously, each sGD carries a recoded cDNA 3’ sequences that restore production of a native protein upon successful integration. However, if accurate integration is not achieved and the sGD element fails to be inherited, NHEJ events can induce lof alleles that will incur severe fitness costs (e.g., lethality or sterility) when either homozygous or when subject to dominantly-acting somatic lethal/sterile mosaicism (ref, SuppFig 1). By selecting gRNAs that cut sequences encoding critical amino acids (e.g., catalytic centers or membrane-tethering sequences), generation of functional NHEJ events can be greatly limited or excluded.

**Fig 1.**
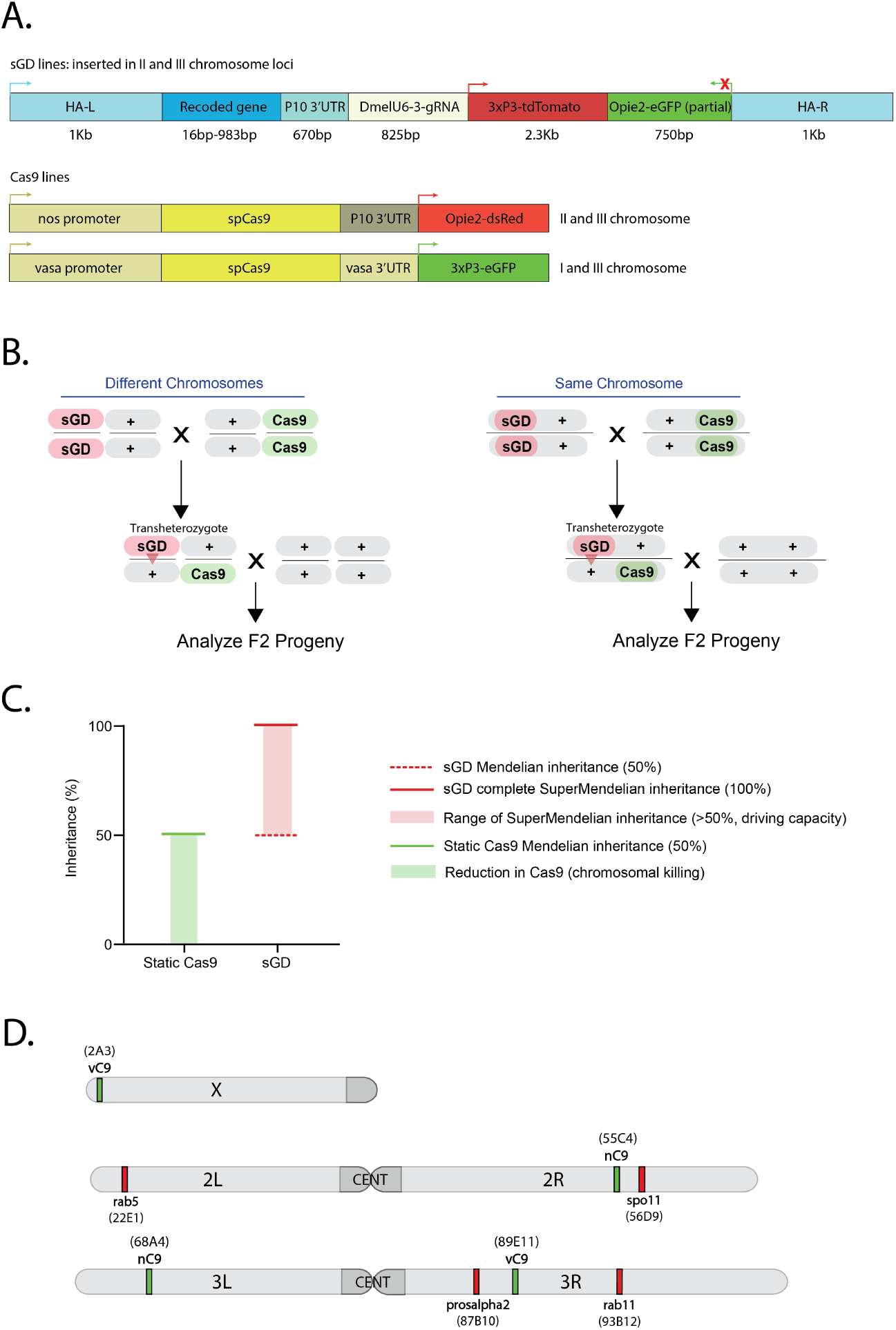
Experimental design of the split gene drive system in recoded essential loci. **A)** Schematic of the genetic constructs engineered and tested in the study. All constructs contain a recoded cDNA of the inserted gene that restores its functionality upon insertion of the transgene, a specific gRNA, and expression of tdTomato driven by 3xP3 to track copying events through fluorescence in the eyes. Static Cas9 lines encode a nanos or vasa-driven Cas9 and a selectable marker, Opie2-dsRed or 3xP3-GFP, respectively. **B)** Outline of the genetic cross schemes used to demonstrate the driving efficiency of each sGD, comparing systems where the sGD, Cas9 and wild-type (+) are located in the same (right panel) or different chromosomes (left panel). F_1_ trans-heterozygotes (carriers of both Cas9 and sGD *in trans*) were singly crossed to wild-type individuals to assess germline transmission rates by scoring % of the fluorescence markers in F_2_ progeny. The conversion event at the sGD locus is shown with a red triangle in F_1_ individuals. **C)** Overview of how data is plotted throughout the paper. F_1_ germline inheritance is plotted in two independent columns, one that refers to the static Cas9, which should be inherited at Mendelian ratios since it does not have driving capacity, and a second column that displays the biased inheritance of the sGD transgene. Graph contains no data. **D)** Chromosomal location and insertion sites of all sGD and static Cas9 transgenes in the *Drosophila melanogaster* genome.

The genes into which we inserted sGDs were all autosomal, located on either the second or third chromosomes (Fig. 1C). All sGD elements targeted conserved DNA coding sequences of recessive lethal (*rab5, rab11, prosalpha2*) or sterile (*mei-W68* [30], the *Drosophila melanogaster* ortholog of *spo11* - referred from here on as *spo11*) genes. These loci were selected based on various criteria including: function and evolutionary conservation (all loci); propitious localization features (membrane-tethered *rab5, rab11* [31, 32]); or stringent cellular requirements (*prosalpha2* [33]). If possible, sgRNA targeted locations near the carboxy-terminus to minimize the extent of recoding necessary (e.g., only the terminal amino acids for rab5 and rab11). Loss-of-function mutants for any of these genes result in different levels of lethality or sterility (*spo11*). For the Cas9 component, we used several static *Streptococcus pyogenes* Cas9 sources, inserted in different chromosomes and driven by different promoters (*vasa* - vCas9 or *nanos* - nCas9), to evaluate the efficiency each sGD.

### sGDs display Super-Mendelian inheritance

We first tested whether the different sGDs would efficiently copy themselves onto a WT chromosome when combined with a static source of Cas9 provided in-trans from the X, II or III chromosomes (Fig. 1B). F_0_ sGD homozygous males or virgin females were crossed to Cas9-bearing lines to obtain trans-heterozygous sGD/Cas9 F_1_ individuals, which we refer to as ‘master females’ or ‘master males’. These master males or virgin master females were then single-pair mated to WT individuals of the opposite sex, and the resulting F_2_ progeny were assessed for sGD copying efficiency by scoring the prevalence of fluorescent markers: red-eye (3xP3-tdTomato) for the sGD element, and green-eye (3XP3-eGFP) for vCas9 or red-body (Opie2-DsRed) for nCas9.

In all of these single-generation crosses, F_2_ progeny displayed Super-Mendelian inheritance of the sGDs ranging from 64.8% to 100% (i.e., >50% as expected by Mendelian transmission), depending on the insertion locus and F_1_ trans-heterozygote sex (Fig. 2). Since these sGDs are inserted in autosomes, we were able to use either sex as the sGD or Cas9 providers in the G_0_ generation. We observed no differential effects between Cas9 being carried by G_0_ males versus females. Interestingly, however, for both elements inserted on II-chromosome genes (2L; *rab5* and 2R; *spo11*), we observed F_1_ sex-dependent differences in transmission of the sGD to F_2_ progeny. When transmitted through F_1_ master females, the rab5 sGD was transmitted at a rate of 96% (±0.8; Fig. 2, SuppFig 2A) for all sGD-Cas9 combinations, while the spo11 sGD transmission was somewhat lower (83%±2.9; Fig. 2, SuppFig 2B). In contrast, when transmitted from F_1_ master males, inheritance of the rab5 sGD dropped to an average of 76% (±3; Fig. 2, SuppFig 2A) and to 65% (±2.1) for the spo11 sGD (Fig. 2, SuppFig 2B), both representing significant decreases (~15%) in transmission efficiencies when conversion occurred in males compared to females (rab5 sGD: *U*=1010, n_1_=197, n_2_=116, p<0.0001; spo11 sGD: *U*=2613, n_1_=145, n_2_=109, p<0.0001). For sGD elements located on the III chromosome (3R; *prosalpha2* and *rab11*), transmission frequencies in F_2_ progeny (tdTomato+) derived from trans-heterozygote F_1_ master females revealed highly-biased inheritance of the sGD, with an average above 98% for *rab11* (Fig. 2, SuppFig 2C) and 99% for *prosalpha2* (Fig. 2, SuppFig 2D) with all Cas9 combinations, with little variation. In the case of the rab11 sGD, transmission from F_1_ master males paralleled the trend seen with the II-chromosome insertions, and was reduced to 85% (±6.5; Fig. 2, SuppFig 2C) (rab11 sGD: *U*=1052, n_1_=154, n_2_=122, p<0.0001). In the case of the prosalpha2 sGD, however, drive efficiency was not appreciably reduced when transmitted by F_1_ master (97.8%±1.3; Fig. 2), well above all other tested F_1_ master male sGD/Cas9 combinations (SuppFig 2D - differences were not statistically significant: prosalpha2 sGD: *U*=7640, n_1_=143, n_2_=113, p=0.256).

**Fig 2.**
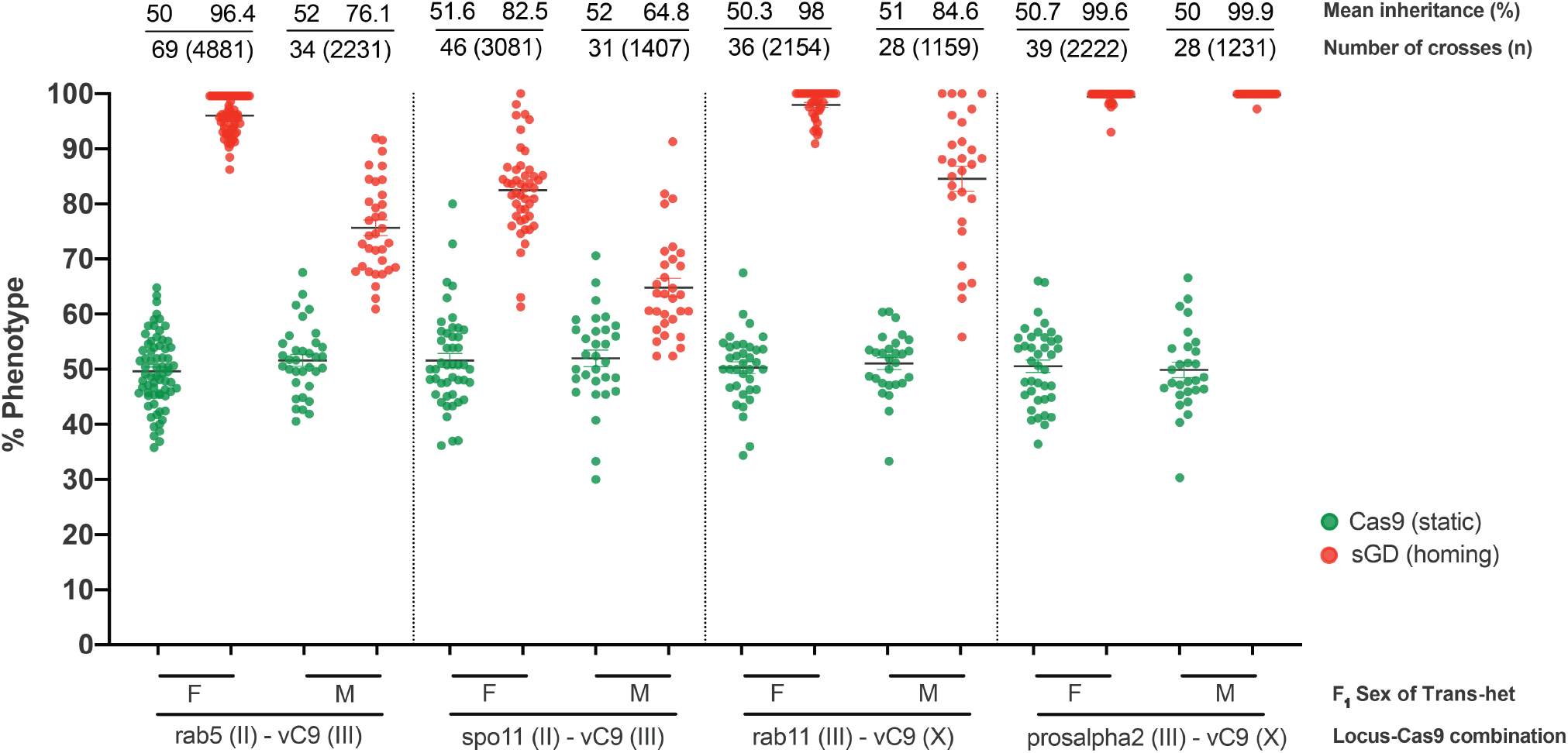
sGD elements display different Super-Mendelian inheritance patterns depending on the transheterozygote progenitor sex. Genetic crosses performed using the sGD transgenes in combination with *vasa*-driven Cas9 lines located in the first (X) or third (III) chromosome. Single F_1_ germline conversion was assessed by scoring the markers for both transgenes in the F_2_ progeny. Inheritance of Cas9 and sGD is depicted using green and red dots, respectively. Each single pair mating cross is shown as a single data point in the Cas9 and sGD columns. Values for the inheritance mean, number of crosses (N) and individuals scored (n) are shown atop of the graph in line with each respective dataset. Sex of the F_1_ parental trans-heterozygote is indicated in the X-axis. Super-Mendelian inheritance is seen for all sGD transgenes. Notably, sGD conversion in crosses from F_1_ females is >95% for three of the four genes tested. However, in crosses derived from F_1_ males, we observed a 15-20% reduction of the transmission observed in F_1_ females for all genes except sGD prosalpha2, where there are no detectable differences depending on the sex of the F_1_ transheterozygote.

Several observations and trends can be derived from the data summarized above. First, locus-specific variation in copying efficiencies that we observed most likely depends on several factors including the genomic context of the insertion locus, gRNA sequence, and the severity of lethal mosaicism (SuppFig 1). Second, the essential nature of the target gene is highly relevant in this context: genes that are broadly required in the organisms for viability (rab5, rab11, prosalpha2) may display varying degrees of lethal mosaicism, depending on whether their activities are required in many or few cells or in a cell-autonomous fashion. These results suggest that stringent requirement for locus function results in more complete and immediate elimination of lof alleles (prosalpha>rab5), whereas for less-vital genes (rab11) some individuals carrying lof alleles are able to survive. Targets that are essential for reproduction, such as spo11, will generate sterile mosaic progeny that survive but are infertile, and most NHEJ events will not be automatically eliminated from the population, but will unable to reproduce efficiently.

Another prominent trend observed in single generation crosses was sex-dependence of sGD transmission, which was observed with all sources of Cas9 and sGDs except for the prosalpha2 drive (SuppFig 3). The basis for sex differences in sGD transmission remains unknown, but may be related to the lack of male recombination in *Drosophila*, which is an unusual genetic characteristic of this species. The prosalpha2 sGD is an interesting exception to this pattern. The absence of a sex-bias for transmission of this drive was observed for both X and 3^rd^ chromosome sources of Cas9 (SuppFig 2D). In prosalpha2 sGD crosses with the 3^rd^ chromosome vCas9-III source, the marker used to score for presence of Cas9 also serves as receiver chromosome marker due to its close linkage to the sGD insertion site (Fig. 1C). In all the other cases, chromosomal distance between the Cas9 source and the sGD does not permit independent tracking chromosomal homologs due to independent assortment. Allele-specific analysis in the case of the vCas9-III source revealed that the receiver chromosome was being transmitted at significantly lower rates than expected (35-40% versus expected 50%). This observation suggests that the copying process sometimes damaged the receiver chromosome resulting in the homolog being inadequately repaired and lost prior to fertilization or shortly thereafter. In crosses of the prosalpha2 sGD to Cas9 sources located on the second or the X chromosomes, we did not observe such deficits in GFP inheritance, consistent with observations that single-cut drive elements generally do not create significant chromosome damage [23]. However, lower levels of chromosome damage may be incurred by unmarked receiver chromosomes, which are difficult to distinguish above background variation in single-generation experiments.

We also evaluated the sGD systems for the phenomenon of ‘shadow drive’, in which maternally-deposited Cas9 in the egg biases the inheritance of a transgene for one extra generation even if the Cas9 transgene is not transmitted to F_2_ females [16, 23]. We crossed F_2_ sGD+/Cas9-virgin females (that only carried maternally-deposited Cas9 protein) to wildtype males and scored for presence of the eye-tdTomato phenotypic marker in F_3_ individuals. In the absence of shadow drive, F_3_ progeny should display Mendelian inheritance of the marker. However, progeny of sGD+/Cas9-F_2_ females exhibited modest levels of super-Mendelian transmission of sGDs, which was most prevalent for rab5 sGD (64.8±11.9% = ~30% conversion, SuppFig 4). We also performed the reciprocal cross of F_2_ sGD+/Cas9-males to wildtype virgin females. As there is no paternal deposition of Cas9 protein via the sperm in such F_2_ males, they transmit the sGDs to their F_3_ progeny, as expected, in at Mendelian rates (~50%, = 0% conversion) (SuppFig 4).

### Resistant alleles do not impede sGD performance

Because lethal mosaicism is hypothesized to act dominantly to eliminate lof NHEJ mutations [23], we analyzed production of these alleles in single generation crosses as well as their predicted elimination in cage studies. Using single-cross experiments (Fig 2), we recovered individual non-fluorescent male and female F_2_ progeny of sGDs inserted into genes essential for viability (i.e., flies lacking the sGD dominant marker) that carried potential NHEJ alleles from one parent and a WT allele on the homologous chromosome provided by the other parent. Knowing the WT sequence, we were able to separate the Sanger sequencing chromatogram reads between WT sequences and those potentially carrying indels. This analysis revealed the production of indels for all genes (including lof frameshift and in-frame mutants) as well as intact WT alleles (Fig 3). For the rab5 sGD, in-frame NHEJ alleles accounted for 34.5% of all NHEJ events (Fig 3A), equating to 1.2% of all individuals. Rab proteins contain carboxy-terminal prenylation sites [34], consisting of two cysteine residues that form disulfide bonds essential for their localization to vesicular membranes. Some in-frame NHEJ events mutated one or both of these two cysteines since, by design, the gRNA cleavage sites were chosen to be very near these codons. Consistent with the role of Rab protein tail residues in forming essential disulfide bonds, none of the cysteine-altering alleles were viable in a homozygous state. 31% of the individuals carried frameshift mutations, which are likely to lead to a protein unable to localize to vesicle membranes and thus defective. For spo11 and rab11, similar categories of mutant alleles were recovered. The spo11 sGD generated few frameshift mutations (Fig 3B) and in-frame mutations accounted for 7.7% out of the total recipient alleles. Since the gRNA cut site for this drive targets a sequence adjacent to critical catalytic residues, many of these in-frame mutants are likely to be sterile when homozygous, as would frameshift mutations leading to truncated non-functional proteins [30]. Rab11 has a similar prenylated 3’ end structure to rab5 [35], with a double cysteine located at the tail end of the protein. Interestingly, no mutants altering these cysteine residues were observed for the sGD in single crosses, even in a heterozygous condition. However, a high percentage (63%, Fig 3C) of the non-fluorescent individuals harbored in-frame NHEJ alleles, accounting for 1.2% of the overall target alleles, similar to the total number seen for rab5 sGD. This marked skew in recovering predominantly in-frame alleles is consistent with lof frameshift alleles being created and eliminated by lethal mosaicism, and validates the strategy of selecting gRNA cut sites in functionally critical domains. In the case of the prosalpha2 sGD, we did not observe a single unaltered WT allele (Fig 3D) among non-fluorescent F_2_ progeny, suggesting Cas9-mediated ~100% cleavage at this locus. This very low rate of total NHEJ events and particularly the lack of in-frame indels (40% of the NHEJ, 0.16% of the total alleles) correlates with the high transmission frequencies observed in single pair crosses and suggests that individuals carrying NHEJ-induced lof alleles rarely survive and/or that the great majority of cleavage events are resolved as copying or chromosomally destructive events.

**Fig 3.**
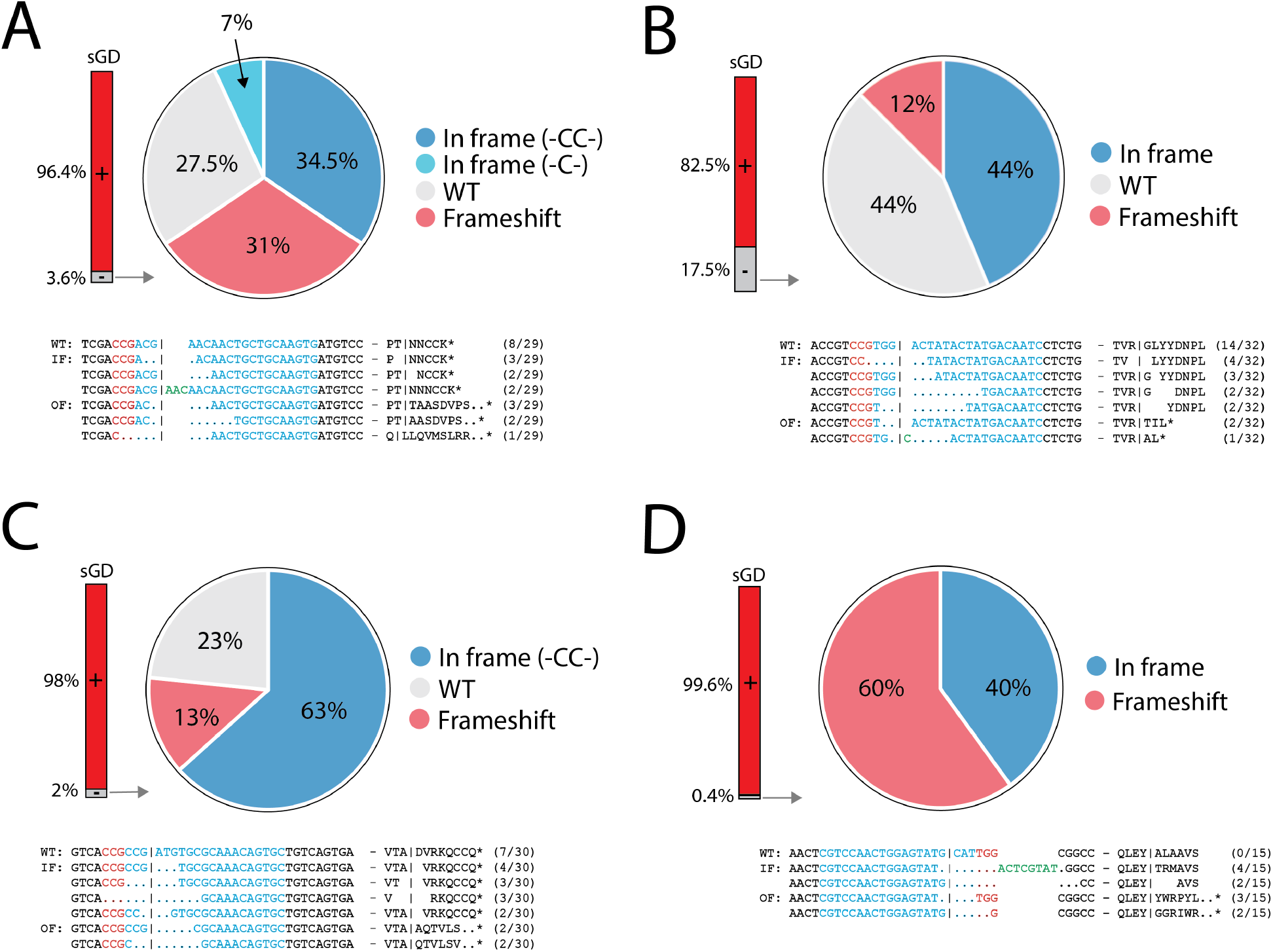
Generation of NHEJ events in single crosses. Sequences were recovered from single non-fluorescent F_2_ individuals generated by the single cross experiments shown in Fig 2. A bar depicts the % of sGD+ (red) and % of non-fluoresenct (sGD-, grey) flies for each tested locus. Genotypic data is depicted in pie charts representing the prevalence of specific indel mutations in sGD-individuals. **A)** rab5, **B)** spo11, **C)** rab11, **D)** prosalpha2. Each section describes the kind of NHEJ that is formed and its percentage among the total tested sGD-(NHEJ/WT) heterozygotes. The specific sequence of prominent NHEJ events, along with its corresponding protein sequence and prevalence, is reported under each pie chart. gRNA sequence of each sGD is depicted in blue and its PAM sequence shown in red.

### sGD exhibit differing degrees of drive and Cas9 depletion in multi-generational cages

To assess the ability of our engineered sGDs to mediate population modification, we tested the performance of the different sGD elements in small laboratory population cages. Generation zero (G_0_) drive-competent master males and master female virgins heterozygous for both the sGD and vCas9 elements (sGD/+;vCas9/+) were combined with WT flies (males and virgin females) at a starting ratio of 1:3 sGD/+; vCas9/+ to WT. Since the drive-bearing flies were heterozygous for the sGD, this initial introduction corresponds to 12.5% drive and Cas9 alleles. Except for the prosalpha2 trials in which the III-chromosome source of vCas9 is closely linked to the sGD insertion locus, the sGD and Cas9 elements were unlinked and could segregate independently. At each generation we scored half of the progeny in a cage for the prevalence of both the tdTomato+ sGD and eGFP+ vCas9 elements based on their fluorescence phenotypes.

The rab5 sGD (Fig 4A) and spo11 sGD (Fig 4B) cage trials were conducted for 20 generations using vCas9-III source. Both of these drives achieved ≥90% introduction by G_5-7_, and remained stable with little variation in overall population sizes or bias in male to female ratios over the remaining generations (SuppFig 5). The long plateau phases observed in all replicates, lasting until G_20_, most likely reflect an equilibrium being attained between the sGD and a small percentage of drive-resistant functional NHEJ alleles. Importantly, gradual progressive reduction in the prevalence of the Cas9 transgene was observed in many cage replicates, which was particularly evident for the spo11 sGD, and may have contributed to the observed multigenerational dynamics. In the rab5 sGD experiments, Cas9 prevalence decreased with variable kinetics among cage replicates, whereas similar slopes of Cas9 decrease were observed in all of the spo11 sGD cage trials. It is notable that following the disappearance of Cas9, whenever it occurred, the prevalence of the rab5 and spo11 sGDs remained stable in subsequent generations (Fig 4A and 4B), indicating that these recoded elements bore no substantial fitness costs relative to other alleles present in the cages (either functional NHEJ or WT alleles).

**Fig 4.**
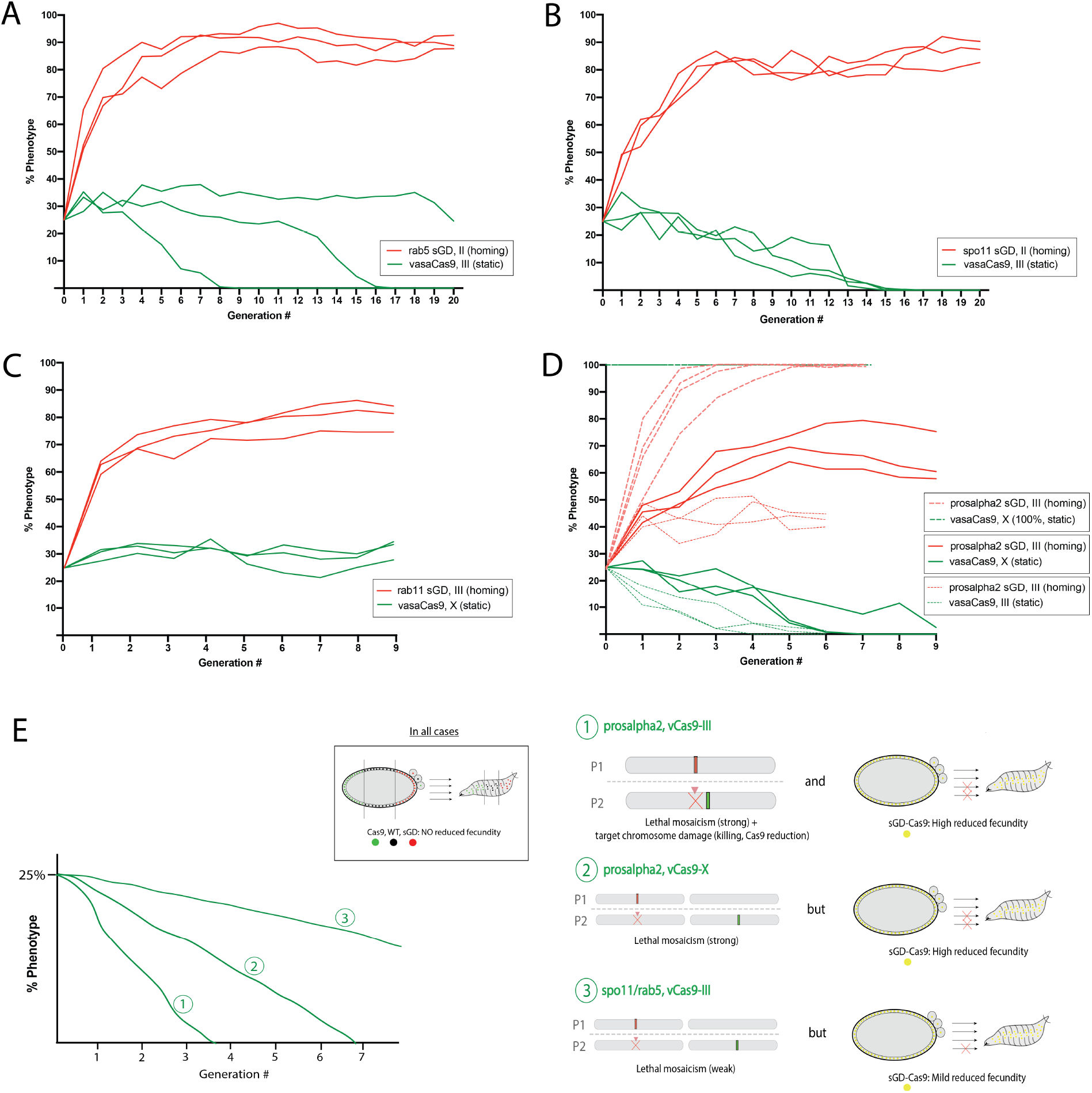
sGD driving experiments in cage trials. Virgin sGD/Cas9 trans-heterozygotes and wild-type flies were seeded at a ratio of 25%:75%. At each generation, flies in a cage were randomly divided in half. One half was scored for eye fluorescence (Generation n) while the other was used to seed fresh cages (Generation n+1). Red traces indicate sGD+ progeny, green traces indicate Cas9+ progeny. Experiments were done in triplicate and each line represents a separate cage. **A)** rab5;vasaCas9(III). The sGD prevalence increases exponentially in the cage (84±6%) up to generation 4 and then plateaus slowly. sGD highest percentage occurs at generation 10 (92±4%). Cas9 decreases in two of the three cages from 25% to 0% by generation 8 and 15, respectively. **B)** spo11;vasaCas9(III). All three replicates reach their highest prevalence in the cage (85±2%) by generation 6-7 and then plateau. Cas9 decreases linearly from 25% to 0% by generation 15 in all three replicates. **C)** rab11;vasaCas9(X). sGD proportions in the cage slowly increase linearly (83±6%) up to the current generation 8. Cas9 does remain at seeding levels (28±3%), suggesting continuous Mendelian transmission. **D)** prosalpha2 sGD drive dynamics depend on the location of the static Cas9, as well as seeding ratios. Bold lines reflect sGD-vasaCas9(X) and dashed lines highlight other sGD-Cas9 combinations. Thin dashes show vasaCas9(III) cage trials and thicker dashes depict drive in a Cas9-saturated population. sGD-Cas9 combinations produce different driving fates and outcomes, providing a flexible tool for deployment. **E)** Hypothesis on cage and drive behavior of the different sGDs.

The drive dynamics for sGD rab11 cages differed somewhat from those of the rab5 and spo11 sGDs in that accompanying a more gradual increase to 80% prevalence by G_8_, the X-linked vCas9-X inserted did not exhibit any obvious decline in frequency (Fig 4C). The higher rates of generating in-frame NHEJ alleles for this the rab11 sGD compared to the rab5 or spo11 sGDs may account for its reduced level of maximal introduction, even with stable maintained inheritance of the vCas9-X transgene (Fig 3C).

Because we hypothesized that lethal (rab5 sGD) or sterile (spo11 sGD) mosaicism may contribute to the drive dynamics, we determined rates of egg laying and hatching for the different sGDs±Cas9. All percentages were normalized to the hatchability observed on WT crosses. We observed reduced hatchability for trans-heterozygote rab5 sGD/Cas9 (84%) and spo11 sGD/vCas9 (91.4%) crosses relative to both sGD/+ (95.5%), vCas9-III (100.5%) crosses (SuppFig 6). The higher hatching rates of the spo11 sGD drive compared to the rab5 sGD may reflect the differential effects of lethal versus sterile mosaicism (see discussion).

### sGD prosalpha2 shows different driving behaviors depending on location of the driving Cas9

As described above, the rab5 and spo11 sGD elements displayed drive trajectories in cage experiments consistent with their performance in single generation crosses. These sGDs drove with similar kinetics and achieved comparable levels of stable introduction on par with their copying efficiencies (considering also that the spo11 sGD is expected to benefit from sterile mosaicism a generation later than rab5 sGD will profit from lethal mosaicism). In contrast, cage trials for the prosalpha2 sGD/vCas9-III displayed a surprising trajectory in which the drive leveled off much sooner than for the other drives (Fig 4D, thin dashed lines). This population behavior contrasts with its high transmission efficiency through both males and females in single generation crosses with the same vCas9 source (>99%). Another notable feature of all three cage replicates was a very rapid decline of the Cas9 transgene (lost in all cages by generation 6). We further analyzed the drive performance of the prosalpha2 sGD using a different source and seeding percentages of Cas9. An X-linked vCas9-X source also declined over time when combined with the prosalpha2 sGD, although more gradually than with its third chromosome vCas9-III counterpart. Accordingly, the prosalpha2 sGD reached higher levels of introduction with vCas9-X (60-70%) than it did with the vCas9-III source (Fig 4D, bold lines), yet was still well below that achieved by the rab5 sGD or spo11 sGD. The decline in the X-linked Cas9 source in these experiments contrasts with stable maintenance of this same element in combination with sGD rab11 in cages, and with its Mendelian transmission in single generation crosses with the prosalpha2 sGD. We also examined hatching rates for this drive, which revealed a significant reduction in embryo viability compared to heterozygotes prosalpha2 sGD/+ (99.2%), vCas9-X/+ (99.3%) (SuppFig 6). Prosalpha2 sGD/vCas9’s hatchability decreased to 79.8% compared to WT crosses. This substantial reduction in fecundity is likely to contribute to the observed cage dynamics, particularly when acting over several generations in randomly mating populations.

Since prosalpha2 sGD was the most promising sGD candidate in single generation crosses due to its ability to achieve >99% transmission (Fig 2), we wondered whether we could increase its level of introgression in population drive experiments by forcing maintenance of Cas9 in the population. To do this, we seeded cages at the same 25% trans-heterozygous sGD;Cas9 rate as in the other cage experiments, but replaced the WT population with vCas9-X homozygous flies and only released trans-heterozygote males (which carry only one copy of vCas9-X), thereby increasing the Cas9 frequency to 100%. In this case, the prosalpha2 sGD drove rapidly to complete fixation (in 3 generations in 3 of the 4 cages, with a 2 generation delay in the 4^th^ cage - Fig 4D, thick dashed lines). Consistent with this highly efficient drive trajectory, no NHEJ events were recovered from the cages analyzed for these events. The highly divergent drive outcomes under these differing scenarios suggests a hypothesis for behavior of the prosalpha2 sGD (Fig. 4E), which also may pertain in a less potent mode to other sGDs, as discussed below.

## Discussion

Genetic modification of insect populations to eradicate disease or reduce crop pest effects has been gaining momentum in recent years, partially propelled by the recent development and optimization of CRISPR-Cas9 gene-drive systems capable of dramatic population transformation [2, 3, 5, 14, 15]. Despite the potential of these systems, the creation of Cas9-cleavage resistant-alleles that impede drive and issues surrounding the potential confinability of self-propagating transgenes driven into native insect populations have been raised as principal biological [9, 10] and regulatory [27, 28, 36] concerns for testing and ultimately deploying such systems. Here, using *Drosophila melanogaster* as a model, we demonstrate that it is possible to partially address these concerns by engineering a range of inherently confinable split-drives targeting essential recessive genes (sGDs). These sGD elements carry recoded target gene sequences that both mitigate the formation of cleavage-resistant alleles and also inherently lead to the loss of separately encoded Cas9 sources in population experiments. Specifically, the elimination of cleavage-resistant alleles relies on lof NHEJ events being gradually culled out of the population by standard Mendelian homozygosis or rapidly eliminated after NHEJ alleles are created by dominantly acting maternal lethal-sterile mosaicism [2, 23]. The rapid loss of Cas9 transgenes appears to depend on two putative mechanisms: damage to the Cas9 chromosome (if it closely linked to the drive on the same chromosome) combined with selection against heterozygotes carrying both the sGD and Cas9 (partial haplo-insufficiency).

### Single generation crosses of sGDs reveal importance of locus context

All sGDs tested in single generation crosses showed Super-Mendelian inheritance ranging from 64.8% up to nearly 100%. One clear trend from these experiments was that all sGDs (except prosalpha2) exhibited significantly higher inheritance rates (~15%) when transmitted by F_1_ master females than by F_1_ master males. Since many components of the HDR DNA repair pathway are shared between repair of damage-induced DSB and meiotic recombination, these shared features may underlie both the peculiar lack of male recombination in *Drosophila* and the reduced rates of HDR-mediated gene conversion resulting in the divergent sex specific sGD drive frequencies we observed. This phenomenon was also observed in other studies using a full genedrive [12] as well as in a trans-complementing framework [26]. Why this sex difference was not also observed for the prosalpha2 drive will require further analysis to address, but as discussed further below, may be related to a combination of highly-efficient copying and a particularly strong form of lethal mosaicism that eliminates nearly all lof alleles (NHEJs or damaged chromosomes) if they are ever generated. It is also noteworthy that when the prosalpha2 sGD drive was placed in-trans to a vCas9-III source located on the homologous third chromosome at a site tightly linked to the prosalpha2 locus, the Cas9 source was inherited at a rate lower than that predicted by random chromosome assortment. This finding suggests that the target chromosome was sometimes being damaged by an imperfect repair process leading to its loss during or shortly after transmission.

### Population studies demonstrate variable levels of drive and confinement

Mathematical modeling based on the promising performance of sGDs in combination with various Cas9 sources in single-generation crosses suggested that these drives should achieve high levels of invasion in population cages. Indeed, the rab5 and spo11 sGDs demonstrated intermediate levels of drive that should be suitable for a variety of limited deployment purposes in the field. Combinations of second-chromosome sGDs with the third-chromosome vCas9 source resulted in fixation of the transgene at 80-95% of the population, while vCas9-III was gradually lost from the population over time. The basis for the decline in vCas9-III transgene frequency is not known, but most likely reflects one of two mechanisms examined in more detail for the prosalpha2 drive below, namely damage to the target chromosome and/or modest fitness costs associated with individuals carrying both the sGD and Cas9 source. The latter mechanism seems most likely for all drives except the prosalpha2 sGD since we observed minor reductions in egg laying and hatching when the sGD element and Cas9 were carried together (when sGD conversion occurs), but not for each of the separate elements. These results demonstrate that sGDs targeting genes essential for viability or fertility can effectively spread in a super-Mendelian fashion, provided that a recoded version of the gene is carried by the propagating sGD element and that the gRNA site is chosen to target functionally critical elements of the gene so as to generate as few functional NHEJ alleles as possible. Such loci can be used as docking sites for a split form of drive as shown here, as well as a full gene drive system or in other strategies such as integral gene drives [37] or daisy-chain drives [38]. We note that our system was designed to be readily hackable to a full-drive configuration. HACK refers to “homology assisted CRISPR knock-in” [29] and in our case, we used the system in such a way that a transgene would be able to get introduced into another. Thus, a normally static source of Cas9 would be able to ‘hack’ and mobilize into the sGD locus, creating a self-spreading full drive element (gRNA and Cas9 would spread together). Indeed, we have obtained preliminary proof-of-principle data that such hacking can be achieved (for the spo11 sGD), and a comparison between such full-drive and split-drive versions of these recoded drives will be assessed in future studies.

The behavior of the prosalpha2 sGD in population cages did not conform to simple expectations based on its performance in single-generation experiments. We observed different behaviors of the transgene depending on the source of Cas9. When the prosalpha2 sGD was seeded together with an autosomal vCas9-III at a 25% initial frequency, the Cas9 transgene was rapidly eliminated from the population (within 3 generations in 3 of the four cages). This precipitous decline in Cas9 could readily account for the attenuated drive of prosalpha2 sGD, which only achieved only modest levels of introduction (~50%), only double that of its seeding proportion. The rapid decrease in Cas9 levels may reflect the damage incurred by the receiver chromosome, evidence for which was observed in single generation crosses discussed above (SuppFig 2D). As noted above, the vCas9-III source also declined to varying degrees in different cage replicates when combined with either the rab5 or spo11 sGDs. Combining the prosalpha2 sGD with an X-linked Cas9 resulted in increased drive, but fell short of the levels attained by the spo11 and rab5 sGDs, most likely due to the fitness costs of carrying vCas9-X and the sGD, which again resulted in elimination of the vCas9-X element from the population. It is noteworthy that the vCas9-X source remained stable in the presence of the rab11 sGD. In addition, for all sGD/Cas9 combinations where Cas9 levels declined, reductions were also observed in egg laying and/or hatching rates.

Importantly, when maintenance of the vCas9-X source was forced in cage experiments by mixing drive-competent prosalpha2 sGD flies with vCas9-X/vCas9-X homozygous flies, the drive rapidly achieved full introduction as would be predicted from its inherent drive potential. We offer the following hypothesis to account for prosalpha2 sGD cage results (Fig. 4E) and their relevance to performance of the other sGDs. According to this model, the prosalpha2 sGD incurs two types of drive-dependent fitness costs in different scenarios. When crossed to vCas9-III, it both damages the target chromosome (as evident in the single generation crosses) and also induces a significant heterozygous fitness cost, perhaps associated with efficient and penetrant lethal mosaicism, leading to a modest haplo-insufficient phenotype and consequent selection against individuals carrying both Cas9 and the prosalpha2 sGD. These two processes act in concert to reduce Cas9 prevalence rapidly. When the prosalpha2 sGD is combined with the unlinked vCas9-X source, it does not inflict appreciable damage to the Cas9 chromosome, but still generates a sGD/Cas9-dependent fitness cost resulting in a more gradual loss of the Cas9 transgene. This combined sGD/Cas9 induced effect is also manifested by the rab5 and spo11 sGDs, which may have similar origins in creating milder forms of recessive alleles in the target chromosome.

Altogether, these experiments offer important insights into design features of the sGD system that can be exploited to achieve specific levels of spread and confinement of these systems when applied into native populations. For example, modest introgression of the sGD followed by rapid removal of the drive can be achieved by placing the prosalpha2 sGD across from a Cas9 source on the same chromosome near the *prosalpha2* locus. Intermediate levels of drive could also be achieved by varying the intensity of lethal mosaicism or by choosing sterile mosaicism, which delays fitness penalties by one generation. As in the case illustrated by the prosalpha2 sGD, one can also vary the ratio of sGD to Cas9 elements to control drive trajectories and outcomes. Targeting loci such as prosalpha2 could also be combined with delivery of an inefficiently acting gRNA that targets Cas9 to eliminate such drives from a population if they retain any residual Cas9-dependent fitness costs when homozygous. Such systems could serve as updating platforms to sustain allelic drives [23] that bias inheritance of beneficial traits such as susceptibility to insecticides, or to disseminate cargo with desired features such as new rounds of anti-pathogenic molecules to supplement those carried by an original modification drive [39, 40].

## Methods

### Plasmid construction

All plasmids were cloned using standard recombinant DNA techniques. Fragments were amplified using Q5 Hotstart Master Mix (New England Biolabs, Cat. #M0494S) and Gibson assembled with NEBuilder HiFi DNA Assembly Master Mix (New England Biolabs, Cat. # E2621). Plasmids were transformed into NEB 5-alpha chemically-competent *E. coli* (New England Biolabs, Cat. # C2987), isolated and sequenced. Full sequences of plasmids and oligos can be found in Supplemental Material.

### Microinjection of constructs

Plasmids were purified using the PureLink Fast Low-endotoxin Maxi Plasmid Purification kit (ThermoFisher Scientific, Cat. #A35895). All constructs were fully re-sequenced prior to injection. Embryo injections were carried out at Rainbow Transgenic Flies, Inc. (http://www.rainbowgene.com). Each gRNA construct was injected into a vasa-Cas9 expressing line in the 3^rd^ chromosome (Bloomington #51324).

Injected embryos were received as G_0_ larvae, were allowed to emerge and 3-4 females were intercrossed to 3-4 males in different tubes. G_1_ progeny were screened for the eye-tdTomato positive marker that indicates transgene insertion. All transgenic flies that displayed the red marker were then balanced using Sco/CyO (for genes located on the II chromosome) or TM3/TM6 (III chromosome) and kept on a w^1118^ background. Homozygous stocks were kept in absence of any balancer alleles. Correct insertions in homozygous transgenic stocks were confirmed through Sanger sequencing.

### Fly genetics and crosses

Fly stocks were kept and reared on regular cornmeal medium under standard conditions at 20-22°C with a 12-hour day-night cycle. sGD housekeeping and crosses were performed in glass vials in an ACL1 fly room, freezing the flies for 48 hours prior to their discard. To assess each locus’ homing efficiency, we genetically crossed each sGD construct to different Cas9 lines. Since our GOIs are autosomal, individual trans-heterozygote F_1_ males or virgin females were collected for each G_0_ cross and crossed to a wildtype fly of the opposite gender. Single one-on-one crosses were grown at 25°C. Inheritance of both gRNA (>50%) and Cas9 (~50%) were calculated using the resulting F_2_ progeny by scoring the phenotypic markers associated to each transgenic cassette.

### Multi-generational Cage Trials

All population cage experiments were conducted at 25°C with a 12-hour day-night cycle using 250 ml bottles containing standard cornmeal medium. Crosses between flies carrying the gRNA and flies carrying vCas9 (X or III) were carried out to obtain F1 trans-heterozygotes used to seed the initial generation. Wildtype or trans-heterozygote males and virgin females were collected and separately matured for 3-5 days. Cages for all loci were seeded at a phenotypic frequency of 25% gRNA/Cas9 trans-heterozygotes (15 males, 15 females) to 75% w^1118^ (45 males, 45 females). In each generation, flies were allowed to mate and lay eggs for ~72 hours, when parents were removed from the cage (G_n_), and kept for 10 days. Subsequent progeny (G_n+1_) were randomly separated into two pools and scored; one was collected for sequencing analyses while the other was used to seed the following generation. This process of random-sampling and passage was continued for 10-20 generations.

### Molecular analysis of resistant alleles

To extract fly genomic DNA for single fly resistant allele sequence analysis, we used a protocol described previously (Gloor GB et al). After extraction, each sample was diluted 1:3 (sample:H_2_O). 1-2ul of each diluted DNA extraction was used as template for a 25ul PCR reaction that covered the flanking regions of the gRNA cut site, which were used to sequence the alleles. Sanger sequencing in individual flies was performed at Genewiz, Inc. in San Diego, CA.

Oligos used for either single fly or cage trial deep sequencing analyses can be found in Supplementary Information.

### Viability assays

For embryo viability counts, 2 to 3-day old virgin female flies were mated to wild-type males for 24h and 15 females (in triplicate) were placed in egg-collection chambers to lay eggs during a 18h period, then the laying plate was removed and eggs counted. All embryos were counted and kept on an agar surface at 20 °C for 48 h and hatchability of those eggs was calculated at that point by counting the unhatched embryos. In all plates, females laid between 211 and 332 eggs. Each experiment was carried out in triplicate, and the results presented are averages from these three experiments. Embryo survival was normalized to the survival observed in parallel experiments carried out with WT flies.

For adult fly counts, the larvae obtained in each embryo count assay replicate were transferred from egg collection plates to 250-mL bottles containing modified cornmeal medium. All adult flies that emerged from these bottles were counted and the results of the three replicates for each experiment averaged together.

### Figure generation and Statistical analysis

Graphs were generated using Prism 8 (GraphPad Software Inc., San Diego, CA) and potentially modified using Adobe Illustrator CS6 (Adobe Inc., San Jose, CA) to visually fit the rest of the non-data figures featured in the paper.

### Safety measures

All sGD crosses were performed in glass vials in an ACL1 facility, in accordance with the Institutional Biosafety Committee-approved protocol from the University of California San Diego. All vials were frozen for 48 hours prior to autoclaving and discarding the flies.

## Acknowledgements

We would like to thank members of the Bier, Akbari and Gantz laboratories for continuous comments and discussions on the paper. We thank Ting Yang for her contribution on generating reagents, Nikolay P Kandul for insights and providing nCas9 fly lines and Xiang-Ru (Shannon) Xu for her help and comments on figures. OSA, AB, IS were supported in part by funding from a DARPA Safe Genes Program Grant (HR0011-17-2-0047) and NIH grants (R21RAI149161A, R01AI151004, DP2AI152071) awarded to OSA. These studies were supported by NIH grant R01 GM117321, a Paul G. Allen Frontiers Group Distinguished Investigator Award to E. Bier (E.B.) and a gift from the Tata Trusts in India to TIGS-UCSD.

## Author Contributions

GT, ABB, OSA and EB conceptualized the study. GT and ABB contributed to the design of the experiments. GT, ABB and IS cloned the constructs, isolated the transgenic lines and gathered preliminary data. GT performed the experiments and analyzed the data. GT designed artwork. All authors contributed to the writing and approved the final manuscript.

