## Supplementary Figures for "Inherently confinable split-drive systems in *Drosophila*"

**This document contains:**

- Supplementary Figures 1-6

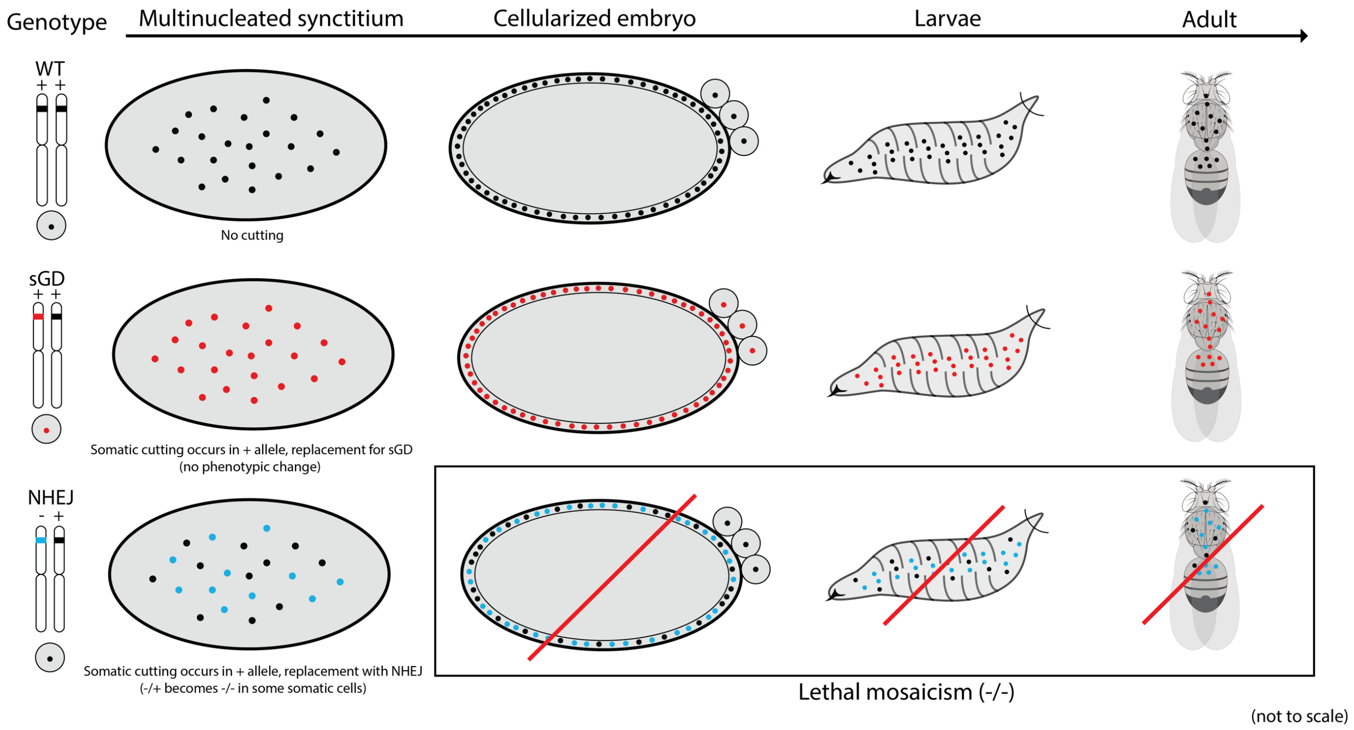


**Supp Fig 1. Schematics of the lethal mosaicism effect.** Scheme explaining the phenomenon of lethal mosaicism. The first two rows (WT, +/+ and sGD, recoded +/+) depict successful fly development to adulthood. The bottom row (NHEJ, +/-) shows the developmental pause following Cas9 cleavage of somatic cells. Cleavage can occur in the parental WT (+) chromosome somatically and generate mutation through repair by NHEJ. This can lead to the generation of a second indel on the homologous chromosome by formation of a new NHEJ event or by copying from the already mutated NHEJ allele. Cells that undergo this somatic conversion will present a knockout phenotype (-/-), leading to cell death when the affected gene is deemed essential for survival. Cellular death and the stage at where the knockout phenotype is visible varies depending on the amount of cells affected, time of cleavage and essentiality of the gene.

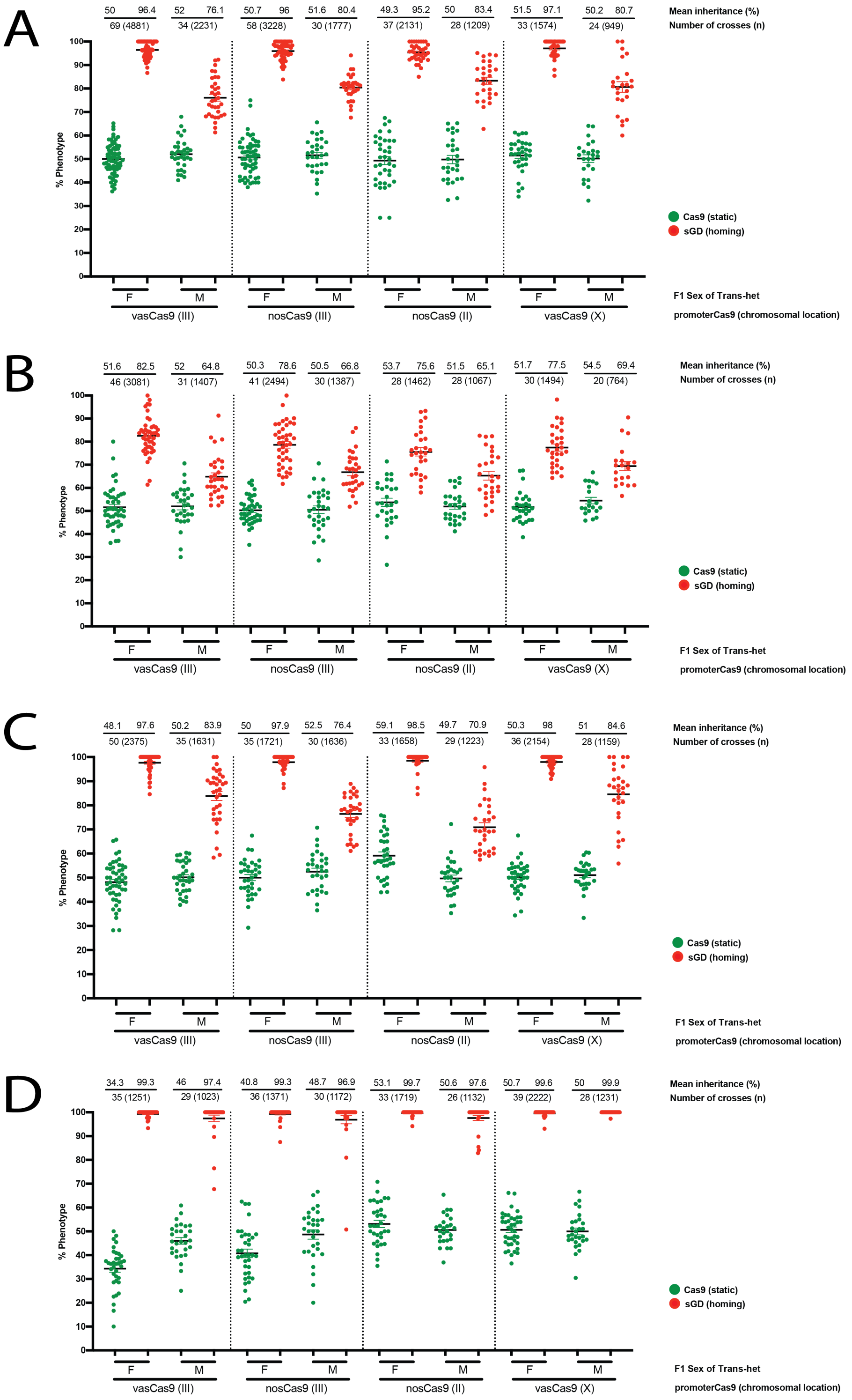


**Supp Fig 2. sGD – Cas9 combinations.** Genetic crosses performed using the sGD transgenes in combination with *vasa* or *nanos*-driven Cas9 lines located in the first (X), second (II) or third (III) chromosome. Graph contains data for **A)** rab5 **B)** spo11 **C)** rab11 **D)** prosalpha2. Single $F_1$ germline conversion was assessed by scoring the markers for both transgenes in the $F_2$ progeny. Inheritance of Cas9 and sGD is depicted using green and red dots, respectively. Each single cross is shown as a single data point. Values for the inheritance mean, number of crosses (N) and individuals scored (n) are shown atop of the graph in line with each respective dataset. Sex of the parental (F1) trans-heterozygote is indicated in the X-axis. Super-Mendelian inheritance is seen across all sGD transgene-Cas9 combinations. However, differences in the germline promoter driving Cas9 and the chromosomal location of the Cas9 transgene can affect the observed inheritance ratios.

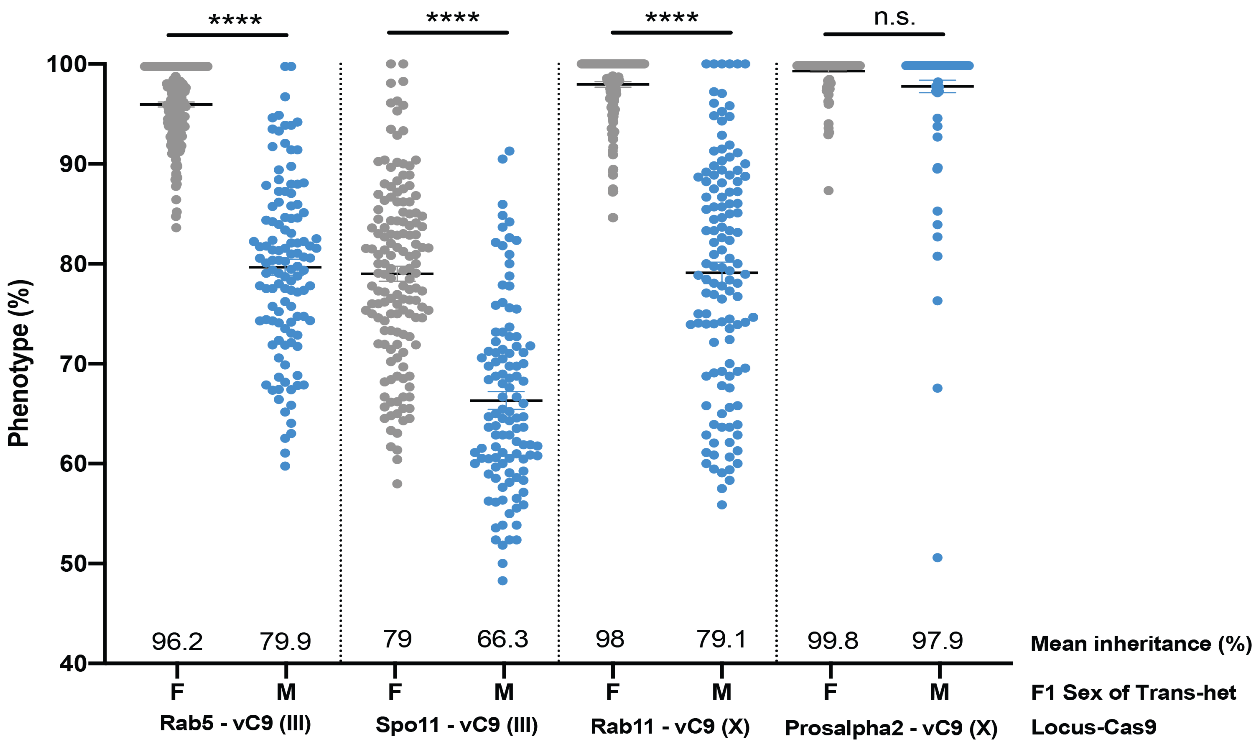


**Supp Fig 3. Pooled inheritance of sGD-Cas9 can be used to observe differences in Super-Mendelian inheritance depending on sex and independent of the Cas9 promoter used.** Graph depicting the differences in $F_2$ inheritance depending on the sex of the $F_1$ trans-heterozygote where germline chromosomal conversion occurs. For all tested genes except prosalpha2, we observed a 15-20% drop in transgene transmission when conversion occurs in $F_1$ males (blue) compared to transmission through $F_1$ females (grey).

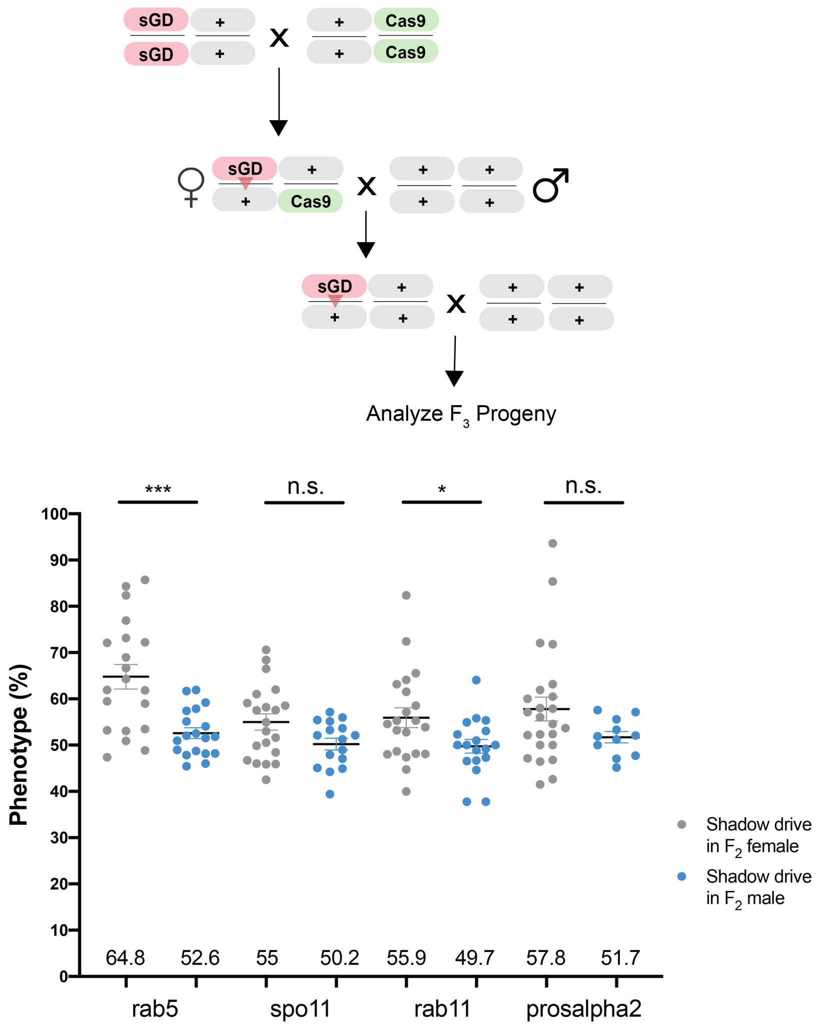


**Supp Fig 4. Shadow drive.** In some instances, maternally-deposited Cas9/gRNA complexes can mediate cleavage of the gRNA target site even in the absence of genetically encoded Cas9 and thus bias transmission rates an extra generation. This phenomenon is known as "shadow drive". To assess the capacity to perform shadow drive for different sGD, we analyzed sGD+/Cas9- $F_2$ individuals, crossing them to a wild-type individual and scoring for the marker phenotype in $F_3$ progeny (shown in cross scheme). In grey, sGD transmission from $F_2$ females; in blue, sGD transmission from $F_2$ males, which serve as a control since there is no paternal deposition of Cas9 into sperm cells.

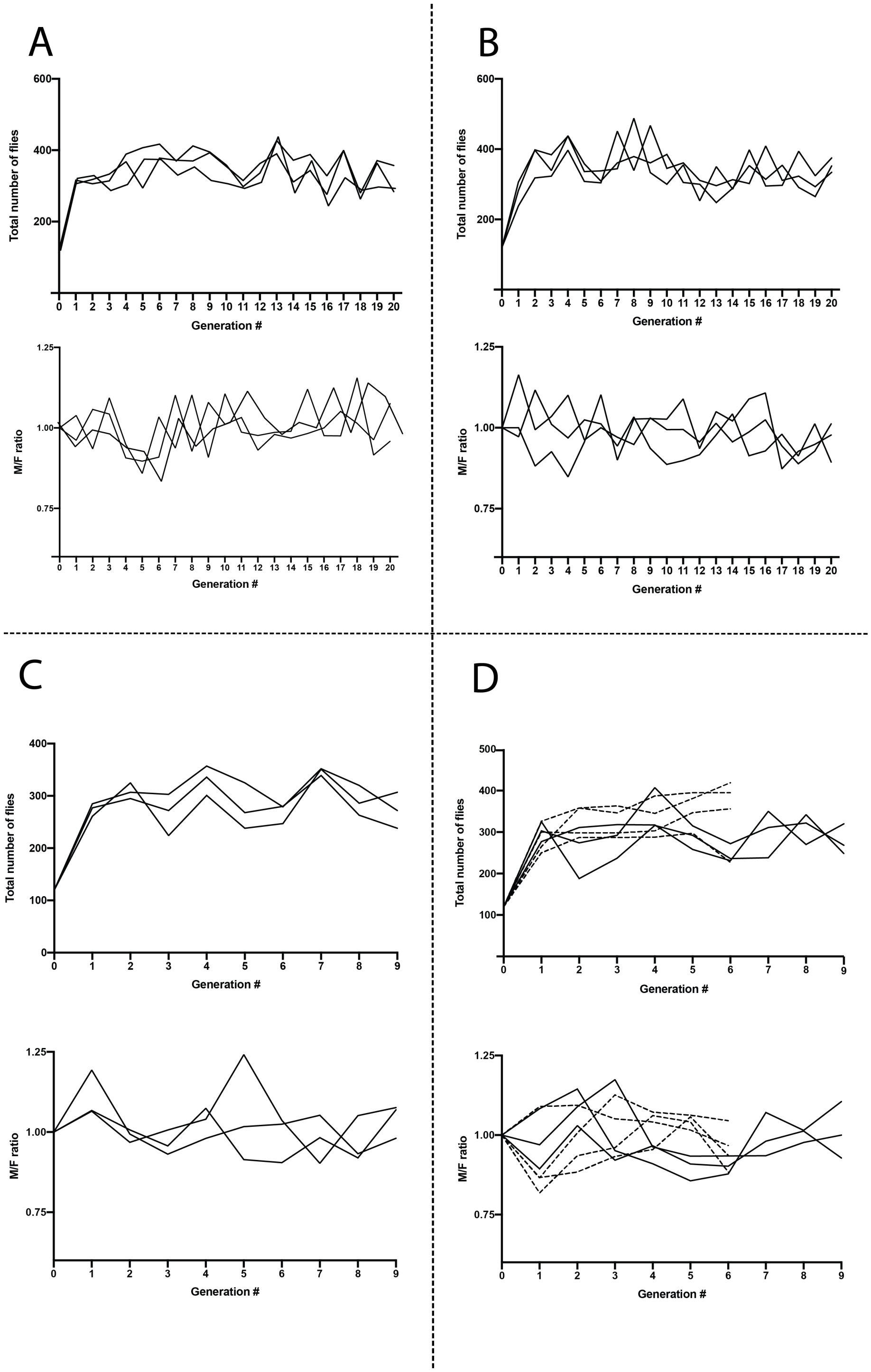


**Supp Fig 5. Population size and male-to-female ratios in cage trials.** Cage trial data for sGD **A)** rab5 **B)** spo11 **C)** rab11 **D)** prosalpha2. Total number of flies and male-to-female ratios were monitored at every generation to unveil possible population size reductions or sex preference due to associated fitness costs. In all conditions, we observed random fluctuations of the population size and sex ratios.

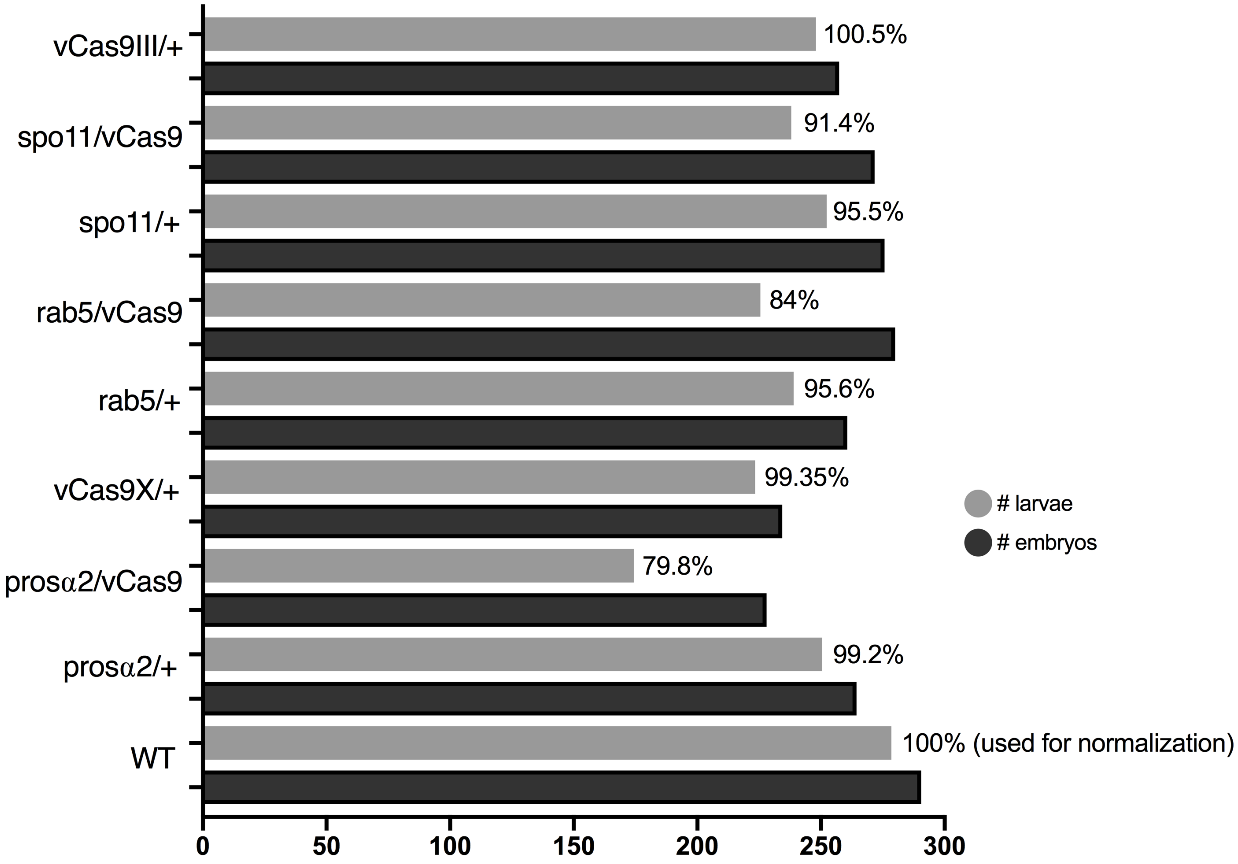


**Supp Fig 6. Egg-laying and hatchability assays.** To account for potential fecundity-associated costs in transgenic flies, we performed a series of egg-deposition and viability experiments. The graph shows number of laid embryos (black) and number of those that hatch (grey). Hatching percentages were normalized to WT fly values. Hatching was significantly impaired in transheterozygotes containing both sGD and vasaCas9, decreasing from >95% to 79.8%, 84% and 91.4% for prosalpha2, rab5 and spo11, respectively. We associate this difference in embryonic viability to somatic lethal mosaicism. Adult fly counts not graphed.
